# Nicotine cessation in female, but not in male mice, mitigates the metabolism-disrupting offspring effect of nicotine exposure

**DOI:** 10.64898/2026.05.24.727521

**Authors:** Amanda Process, Raquel Chamorro-Garcia, Carlos Diaz-Castillo

**Affiliations:** Department of Microbiology and Environmental Toxicology, University of California, Santa Cruz, Santa Cruz, CA

**Keywords:** epigenetics, gamete development, multigenerational metabolism-disrupting agents, nicotine, nicotine aversion, nicotine cessation

## Abstract

**Introduction:** It is now widely recognized that environmental exposures can predispose unexposed descendants to disease across multiple generations without inducing genetic mutations. Among the numerous unknowns that multigenerational effects still hold, identifying the most probable windows of susceptibility for multigenerational environmental disease predisposition remains a crucial challenge in preventing such effects. We have proposed that multigenerational environmental effects can be mediated by perturbations in chromatin organization that originate from environmental exposures causing alterations in gamete elements necessary for establishing chromatin organization immediately after fertilization. Based on this hypothesis, it is likely that the period preceding conception serves as a relevant window of susceptibility for multigenerational effects, and that such susceptibility may vary between female and male preconception exposures due to the distinct characteristics of oocytes and sperm. Here, we test this framework using nicotine—a well-established endocrine- and metabolism-disrupting chemical with documented multigenerational effects—and assess whether windows of nicotine cessation prior to conception that span the last stages of gamete maturation mitigate these effects.

**Methods:** We conducted two asynchronous studies to determine the direct and offspring effects of female preconception exposure (FPE) and male preconception exposure (MPE) to nicotine and nicotine cessation. We exposed C57BL/6J female (FPE) or male (MPE) mice to deionized water (control), continuous nicotine (300 µg/mL), or one of two nicotine cessation windows whose durations did or did not encompass one full round of gamete maturation. Following exposure, we mated exposed mice with unexposed mice of the same age to produce their offspring. We measured the water and food consumption and body weight of exposed mice to determine the efficacy and direct effect of the assayed exposures. We also measured body weight, fasting body weight, fasting glucose, gonadal white adipose tissue and liver weights, and plasma concentrations of twelve metabolic hormones in the offspring of exposed mice to determine the offspring effect of nicotine exposure and its mitigation upon nicotine cessation. We determined the significance of comparisons between nicotine and control groups using the Monte Carlo-Wilcoxon testing framework that we have previously developed.

**Results:** Preconception nicotine exposure elicited sexually dimorphic metabolic effects in the offspring of exposed mice that differed between FPE and MPE studies. Nicotine cessation mitigated F1 metabolic perturbations only after maternal—not paternal—preconception exposure, and only when the cessation window encompassed one full round of oocyte maturation.

**Conclusions:** These findings support the hypothesis that preconception exposures perturb offspring metabolism through sex-specific gamete mechanisms and highlight that the efficacy of cessation strategies depends on the parental sex exposed.

## Introduction

It is no longer a matter of debate that the effect of certain environmental exposures can be transmitted across generations independently of genetic mutations predisposing unexposed descendants of multiple generations to disease (1–3). Although a comprehensive understanding of the mechanisms underlying such phenomena and which current disease predispositions could have been influenced by which ancestral exposures is still under investigation, a highly relevant aspect remains unexplored: identifying the most probable windows of susceptibility for the multigenerational environmental predisposition to disease to help design strategies to prevent such effects in prospective parents.

By employing a murine model to investigate the transgenerational effects of exposure to the biocide tributyltin (TBT), we proposed that environmental exposures can induce perturbations in chromatin organization, which can be transmitted across generations and predispose individuals to metabolic disease (4,5). Building upon such research and the knowledge of the developmental establishment of chromatin organization, we further suggested that preconception is a likely relevant window of susceptibility for environmental exposures to cause perturbations in germ line elements (6,7). These perturbations subsequently disrupt the establishment of chromatin organization in the offspring of exposed individuals, which can be propagated through development and across generations (6,7). We recently found supporting evidence for this hypothesis by examining the effects of exposing prior to conception female mice to a panel of diverse metabolic disruptors and male mice to nicotine (8,9).

Here, we present our initial findings investigating the metabolic disruptions observed in the offspring of female and male mice exposed prior to conception to nicotine, a well-known endocrine- and metabolism-disrupting chemical (10,11). We hypothesize that such disruptions may be mediated by alterations in the germ lines of exposed mice, which subsequently influence chromatin organization in their offspring. While it is reasonable to speculate that environmentally-mediated perturbations in chromatin organization in the offspring of exposed individuals could be attributed to alterations in epigenetic marks directly located on the germline chromatin, such as DNA methylation or histone modifications, or proteins and RNAs in the germ cell cytoplasm, recent literature has provided evidence suggesting that alterations in the physicochemical milieu of the zygote immediately following fertilization can also lead to the formation of chromatin organization perturbations (7,12,13). Although a comprehensive study of all these factors is necessary, we have opted for a simpler, albeit indirect, strategy that is relevant in the context of preventing the nicotine-mediated multigenerational effects.

To determine whether preconception exposure to nicotine results in metabolic disruption of the offspring that can be mediated by germ line alterations, we proceed to compare the direct and offspring effects of exposing mice to nicotine and two windows of nicotine cessation with unexposed controls. The two windows of nicotine cessation were defined based on the period of maturation of one round of mature gametes in female and male mice (14,15). Consequently, the longest and shortest windows encompass and do not encompass, respectively, the maturation of the round of mature gametes that will contribute to the generation of offspring in exposed mice. The results of these analyses would reveal whether nicotine and/or their metabolization products, such as cotinine, could directly or indirectly influence the maturation process of germ cells in individuals exposed to them.

Despite the ongoing decline in the prevalence of nicotine-related products (NRPs) consumption in recent decades, NRP consumption remains a significant public health concern for several reasons. Firstly, the global consumption of nicotine-related products remains substantial, surpassing 1.2 billion consumers in 2024 (16). Secondly, the nicotine-related industry has historically demonstrated its adaptability by identifying new target markets for newer products marketed as safer alternatives. In recent years, combustion-free products based on chemically-synthesized nicotine, such as vapers or nicotine pouches, are marketed as safer because they eliminate exposure to combustion by-products and additives (17–19). Lastly, recent evidence suggests that pervasive health effects of nicotine exposure can be observed in the descendants of nicotine users, even in the absence of their own consumption (20–22). This implies that while the pursuit of a nicotine-free society, as desirable as it may be, would not entirely eradicate the health concerns arising from the excessive consumption of nicotine-related products by previous generations.

After not being exposed to nicotine, as complicated as it is considering the effects observed of second and third-hand exposures (23,24), the cessation of the consumption of nicotine products is the most effective measure to combat the harms of such consumption (25). Despite the accumulating evidence on the multigenerational consequences of nicotine consumption and the paramount importance of nicotine cessation as the most effective strategy to mitigate the pervasive effects of nicotine exposure, the study of the mitigation impact of nicotine cessation on the multigenerational effects of nicotine exposure remains limited (26,27).

## Materials and Methods

### Mouse procedures and sample processing

We conducted all mouse procedures at the University of California, Santa Cruz (UCSC) Vivarium, in compliance with the animal protocol Diazc2204, which had previously been approved by the UCSC Institutional Animal Care and Use Committee (IACUC). We conducted two asynchronous studies with comparable structures, unless otherwise indicated (Figure 1). The female preconception exposure (FPE) study was conducted in 2023, while the male preconception exposure (MPE) study was conducted in 2024. We procured C57BL/6J mice from Jackson Laboratory (strain #000664). For the FPE study, we procured 80 female and 40 male mice, whereas for the MPE study, we procured 80 female and 80 male mice. To facilitate the acclimation of mice to the UCSC Vivarium, we scheduled their arrival at least one week prior to the time they were required. Upon the arrival of the mice to be exposed, we ear-punched each mouse and introduced them into the Chamorro-Garcia group mouse records, assigning them a unique identifier. We weighed each mouse by carefully placing it in an empty beaker atop a standard laboratory balance. To minimize any initial disparities in body weight between the experimental groups, we randomly assigned 4-mouse cages to four distinct groups as previously described (8). Briefly, we ranked the mouse cages based on the cumulative body weights of the mice within each cage, from highest to lowest. Subsequently, we divided the ranked list into five subgroups of four cages each and randomly assigned each cage to an experimental group.

**Figure 1.**
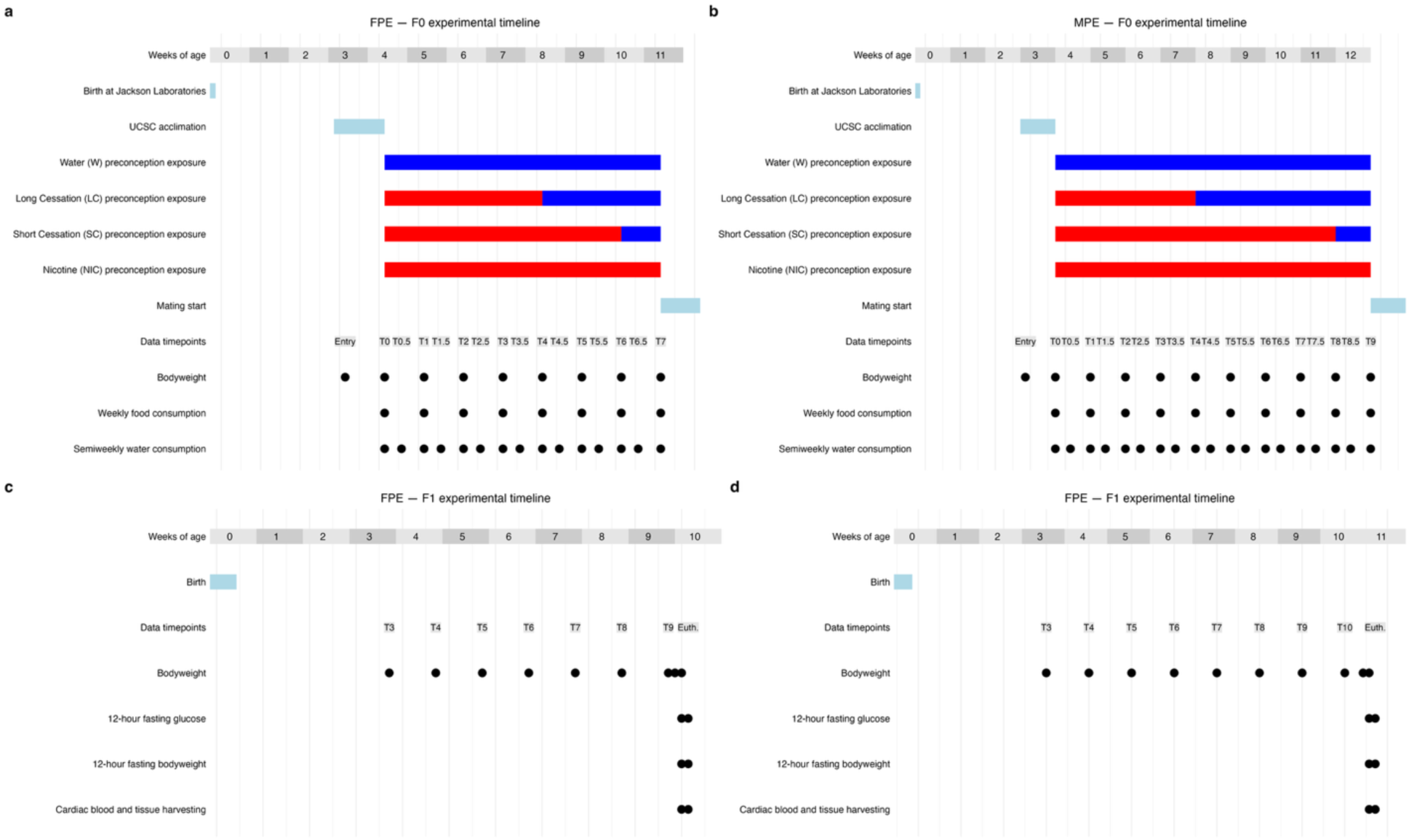
Experimental timeline for the study of female and male preconception exposure (FPE and MPE, respectively) to nicotine and two windows of nicotine cessation (long and short). (**a, b**) Experimental timeline detailing primary mouse operations and the timepoint structure used to assess variations in body weight, water, and food consumption caused by the direct exposure of female **(a)** or male **(b)** mice to nicotine and nicotine cessation when compared to controls. (**c, d**) Experimental timeline detailing primary mouse operations and the timepoint structure used to assess variations in litter size, sex ratio, body weight, fasting body weight, fasting glucose, tissue weight, and plasma metabolite concentrations induced by female **(c)** or male **(d)** preconception exposure to nicotine and nicotine cessation when compared to controls. UCSC: University of California, Santa Cruz; FPE: female preconception exposure; MPE: male preconception exposure; W: water (control group).

Beginning at four weeks of age, we exposed each female mouse group in the FPE study and each male mouse group in the MPE study to four distinct drinking water treatments (Figure 1). We treated the water (W) group in each study, which served as the negative control, with deionized (DI) water for either 7 (FPE) or 9 (MPE) weeks. We treated the nicotine (NIC) group in each study with 300 µg/mL (-)-nicotine (CAS number: 54-11-5; Sigma-Aldrich N3876) in DI water for either 7 (FPE) or 9 (MPE) weeks. We treated the long cessation (LC) group in each study with 300 µg/mL (-)-nicotine in DI water for 4 weeks followed by no-nicotine DI water for either 3 (FPE) or 5 (MPE) more weeks. We treated the short cessation (SC) group in each study with 300 µg/mL (-)-nicotine in DI water for either 6 (FPE) or 8 (MPE) weeks followed by no-nicotine DI water for 1 more week.

To ensure the efficacy of water treatments and to assess the metabolic responses of exposed mice, we conducted repeated measurements of mouse body weight and food and water consumption throughout the exposure period. At the start of the exposure and weekly thereafter, we weighed mice as previously described. At the start of the exposure and semiweekly thereafter, we provided each mouse cage with 200 mL of treated water (entry water) and measured the volume of unconsumed water after three or four days (exit water). At the start of the exposure and weekly thereafter, we supplied mouse cages with approximately 200 grams of fresh mouse food (entry food) and weighed any unconsumed food after each week (exit food).

At the end of the exposure period, we mated the exposed mice with unexposed same-age mice of the opposite sex for a week to produce their F1 offspring. We provided mating cages with unrestricted, untreated mouse feed and drinking water. For the FPE study, we mated two exposed females from the same cage with one unexposed male (2:1 matings). For the MPE study, we mated one exposed male with one unexposed female (1:1 matings). At the conclusion of the mating week, we returned female mice to 4-mouse cages with their same cage mates from the exposure (FPE) or acclimation (MPE) periods for the remainder of their pregnancies. We determined successful matings by monitoring daily for copulatory plugs during the mating week and body weight gains exceeding 1.75 grams observed eight days after the conclusion of the mating week (28). Two days prior to the anticipated birth, we relocated the pregnant females to individual cages equipped with ample bedding material, enabling them to construct a nurturing nest. We closely monitored births and the welfare of litters using the least invasive methods possible until they reached a sufficient age for further manipulations.

At 8-11 days of age, we toe-clipped, counted and sexed all F1 mice and introduced them into the Chamorro-Garcia group mouse records, assigning each one a unique identifier. We used per-litter mouse counts to determine litter size and sex bias as the percentage of female mice. At 3 weeks of age, we weaned F1 mice and selected one female-male sibling pair from 10 litters per exposure group. We housed mice of the same sex and exposure group together in cages with unrestricted, untreated feed and water. We weighed the F1 mice at weaning and continued to do so weekly thereafter.

At 9-11 weeks of age, we weighed F1 mice before and after a 12-hour fasting period. Subsequently, we euthanized mice using an overdose of isoflurane followed by cardiac exsanguination to determine fasting glucose and collect cardiac blood, gonadal white adipose tissue (gWAT) depots, and liver. We performed dissections by all appropriately trained members of the Chamorro-Garcia group. To minimize the daily dissection workload, we euthanized and dissected all female and all male mice in two consecutive days, respectively. The lead experimenter (Diaz-Castillo) employed a randomized approach to assign the order of mouse dissection and assign each mouse to a dissector without prior knowledge of their group.

To determine fasting glucose, we used a Contour® blood glucose meter (BAYER) and Contour® blood glucose strips (BAYER) with a drop of blood drawn from the mouse tails by puncturing them with a needle. To determine plasma levels of relevant metabolites, we drew ∼600 µL of blood directly from the heart using EDTA-flushed syringes to minimize coagulation and collected cardiac blood samples in microcentrifuge tubes containing 6 µL of 100X protease inhibitors cocktail (Sigma Aldrich NC2042678). To separate plasma, we centrifuged blood samples at 5,000 x g for 10 minutes at 4°C. We submitted 50 µL of plasma samples to Eve Technologies (Calgary, Canada) for the determination of the plasma concentration of 12 metabolites included in the Mouse, Rat Metabolic Array (MRDMET): amylin, C-peptide 2 (connecting peptide 2), ghrelin, GIP (gastric inhibitory peptide), GLP-1 (glucagon-like peptide 1), glucagon, insulin, leptin, PP (pancreatic peptide), PYY (peptide Y), resistin, and secretin. Additionally, we weighed gWAT and liver harvested for each mouse using a precision balance.

### Data processing and statistics

We used the following R packages for data processing, result visualization, and figure preparation for publication: *cowplot* (version 1.2) (29), *data.table* (version 1.17) (30), *ggplot2* (version 3.5) (31), *patchwork* (version 1.3) (32), and *scales* (version 1.3) (33). A copy of the R script used for data processing, analysis, and figure preparation for publication will be made publicly available.

For the F0 generation, we compared mouse body weight, water and food consumption between each exposure group (LC, SC, and NIC) and the control group (W) for each exposed sex in each preconception exposure study separately (FPE and MPE). For the F1 generation, we compared litter size and sex bias between each exposure group (LC, SC, and NIC) and the control group (W) for each preconception exposure study separately (FPE and MPE). For the F1 generation, we also compared mouse body weight, fasting body weight, fasting glucose, tissue weight, and plasma metabolite concentrations between each exposure group (LC, SC, and NIC) and the control group (W) for each sex and each preconception exposure study separately (FPE and MPE).

To mitigate potential confounding factors, we normalized body weight for each mouse based on its days of age. Additionally, we normalized fasting glucose, tissue weights, and plasma metabolite concentrations for each mouse relative to its fasting body weight and days of age at euthanasia. We set to zero plasma metabolite concentration data that fell outside the range of the standard curve or that could not be mathematically extrapolated.

We assessed the significance of differences in body weight, litter size, litter sex bias, fasting glucose, tissue weight, and plasma metabolite concentrations for each exposure *versus* control contrast using unmatched-measures Monte Carlo-Wilcoxon (uMCW) tests. We assessed the significance of differences in entry and exit food and water, and of pre-and post-fasting body weights for each exposure versus control contrast using matched-bivariate Monte Carlo-Wilcoxon (mbMCW) tests.

Monte Carlo-Wilcoxon (MCW) tests are particularly well-suited for the integration of analysis of diverse types of data that may follow different distributions, ranges, or scales (8,34). Despite there are four distinct MCW tests to interrogate various data structures, all of them operate similarly and yield comparable metrics that are easily interpretable and comparable (8,34). Briefly, all MCW tests operate by computing a bias index (BI) that spans the range of 1 to −1, indicating that the measure under analysis is completely biased in each conceivable direction. To ascertain the statistical significance of the BIs computed for the user-provided dataset (observed BIs), a series of expected-by-chance BIs are generated by repeatedly rearranging the original dataset and computing BIs for each iteration. *P_upper_* and *P_lower_* values are subsequently calculated as the proportions of expected-by-chance BIs that exhibit values equal to or higher than and equal to or less than the observed BIs, respectively. MCW tests can follow two paths based on the user-defined parameter *max_rearrangements.* If the number of potentially distinctive data rearrangements is less than *max_rearrangements*, MCW tests will draw all potentially distinctive rearrangements, and *P_upper_* and *P_lower_* values will represent an exact estimation of the significance of observed BIs. If the number of potentially distinctive data rearrangements is greater than *max_rearrangements*, MCW tests will perform a number of rearrangements equal to *max_rearrangements*, and *P_upper_* and *P_lower_* values will represent an approximated estimation of the significance of observed BIs. uMCW tests assess whether two sets of unmatched measures (*e.g.*, *a* and *b*) exhibit significant bias in the same direction. mbMCW tests assess whether two sets (*e.g.*, *x* and *y*) of inherently matched-paired measures for two conditions (e.g., *a* and *b*) are significantly differentially biased in the same direction. For all uMCW and mbMCW tests, we used the R package *MCWtests* (version 1.0)(34). We set the *max_rearrangements* parameter to 10,000 and used the same threshold of significance of *P* = 0.05.

We also performed Principal Component Analysis (PCA) to determine the similarities between mouse groups using separately data for F1 female and male plasma metabolite concentrations. We used the function *prcomp* from the R package *stats* (version 4.5) to perform the PCA analyses (35), and the functions *fviz_eig* and *fviz_pca* from R package *Factoextra* (version 1.0) to visualize the results (36). To optimize the value of biological replication, we conducted PCAs utilizing only metabolites whose plasma concentrations had been successfully determined in all biological replicates for all groups in each sex and preconception study. To visually and analytically assess the significance of the separation of mouse groups in PCA plots, we obtained the coordinate matrix utilized for projecting original observations onto the principal component (PC) biplots generated by *prcomp*. We used the coordinate data for the initial two principal components (PC1 and PC2) for each F1 sex and preconception study to generate density plots for PC1 and PC2 that were aligned with each PCA biplot axis (Figure 4e, g, I, k). We also conducted uMCW tests using the coordinate data for PC1 and PC2 separately for each exposure *versus* control contrast.

## Results

### Effects of directly exposing female and male mice to nicotine and nicotine cessation

We conducted two asynchronous studies to determine the offspring effects of exposing C57BL/6J female or male mice prior to conception to nicotine and two windows of nicotine cessation (Figure 1). We randomly divided 80 female mice in the female preconception exposure (FPE) study and 80 male mice in the male preconception exposure (MPE) study into four groups per study. In each study, we exposed each of the four groups to: DI water for either 7 (FPE) or 9 (MPE) weeks (W group; negative control), 300 µg/mL (-)-nicotine in DI water for either 7 (FPE) or 9 (MPE) weeks (NIC group), 300 µg/mL (-)-nicotine in DI water for 4 weeks followed by no-nicotine DI water for either 3 (FPE) or 5 (MPE) more weeks (LC group), and 300 µg/mL (-)-nicotine in DI water for either 6 (FPE) or 8 (MPE) weeks followed by no-nicotine DI water for 1 more week (SC group).

Numerous animal models have been used to investigate the effects of nicotine and its cessation using a disparate array of nicotine exposures (37). In our studies, we opted for a simple nicotine administration method. We exposed mice to nicotine in DI water devoid of any additives that could mask its odor or taste as their sole drinking source. We selected a nicotine exposure concentration of 300 µg/mL because it has been previously demonstrated to elicit plasma levels of cotinine, the primary metabolite of nicotine, indicative of chronic nicotine exposure (38). We established the long and short cessation windows, so they did and did not encompass, respectively, the maturation of one round of mature oocytes (3 weeks in the FPE study) and sperm (5 weeks in the MPE study) (14,15). It has been demonstrated that nicotine and cotinine plasma and urine levels decline to baseline levels after several weeks of nicotine exposure via drinking water as the sole source of water, and only 24 hours of nicotine cessation for male mice (39–41). Therefore, we can argue that although the shortest duration of nicotine cessation in our studies (1 week) would not impede that nicotine and/or its metabolites partially altered the maturation of gametes in exposed individuals, it is unlikely that they are in sufficient amounts in gametes or circulating in the gestating mother to directly impact the development of their offspring. We did not test levels of plasma cotinine to verify our inferences in these studies to avoid disturbing mice in excess, since it is unclear how such types of stresses can contribute to offspring effects.

To assess the efficacy of the treatments and metabolic perturbations in exposed mice, we used a minimally invasive approach that avoided the use of experimental variables that could potentially serve as secondary metabolic disruptors, thereby introducing confounding factors such as fasting. We compared semiweekly water consumption per mouse cage, and weekly food consumption per mouse cage between exposure groups and the control group using mbMCW tests (see Materials and Methods). We also compared mouse body weight between exposure groups and the control group using uMCW tests (see Materials and Methods).

The analysis of water consumption generally revealed that mice treated with nicotine-laced water exhibited a tendency to consume less water compared to mice treated with nicotine-free water (Figure 2a-f). Notably, mice in LC and SC groups showed an increase in water consumption immediately following the cessation of nicotine exposure (Figure 2b-f). These patterns align with nicotine aversion in mice (39,40,42). While nicotine aversion appeared to remain unchanged throughout the nicotine exposure windows for female mice (FPE) (Figure 2b, d, e), it showed a reduction in male mice (MPE) at approximately 3-4 weeks of exposure and further diminished after 8 weeks of exposure (Figure c, d, f). It has been reported that prolonged nicotine exposure mitigates nicotine aversion in male mice (42).

**Figure 2.**
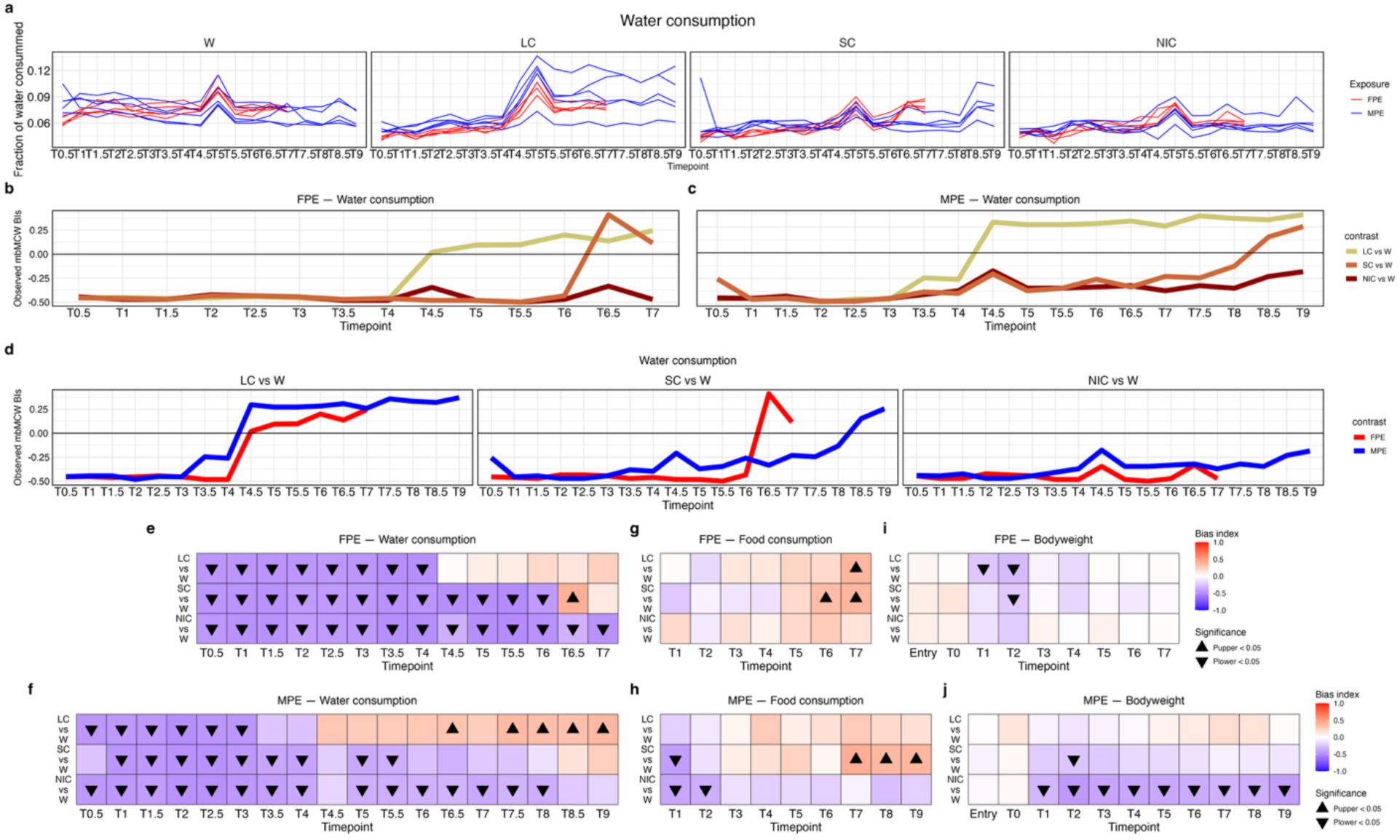
Direct effects of exposing F0 female and male mice to nicotine and nicotine cessation. (**a-h**) Comparison of water consumption (mL) between exposure and control groups using mbMCW tests. (**a**) Water consumption dynamics for each mouse within each experimental group. (**b-d**) Comparison of differences in water consumption between nicotine and nicotine cessation groups when compared with the control group for female and male preconception exposure measured using mbMCW test BIs (see Materials and Methods) shown for each preconception exposure study (**b, c)** or group for each exposure *versus* control contrast (**d**). (**e, f**) Comparison of water consumption (mL) between exposure and control groups using mbMCW tests. (**g, h**) Comparison of food mass consumption between exposure and control groups using mbMCW tests. (**i, j**) Comparison of body weight normalized by age (g/days old) between exposure and control groups using uMCW tests. Sample size for each experimental group is 5 for water and food consumption analyses, and 20 for mouse body weight analyses. mbMCW test: matched-bivariate measures Monte Carlo-Wilcoson test; uMCW test: unmatched measures Monte Carlo-Wilcoson test; BIs: bias index; W: water (negative control), LC: long cessation; SC: short cessation; NIC: nicotine.

The analysis of food consumption revealed comparable trends to the ones observed for water consumption, albeit less pronounced, in nicotine-exposed males (MPE): they exhibited a tendency to consume less food than controls at the onset of nicotine exposure, while simultaneously consuming more food than controls coinciding with the periods when nicotine aversion appeared to abate and with nicotine cessation (Figure 2h). In contrast, the food consumption patterns between nicotine-exposed females (FPE) and control mice did not exhibit significant differences at the beginning of the exposure period (Figure 2g). However, towards the end of the exposure period, female mice in all exposure groups exhibited a tendency to consume more food than controls, with these trends being statistically significant only in the cessation groups (Figure 2g).

The analysis of mouse body weight revealed that the body weight of nicotine-exposed mice decreased when compared with controls immediately at the onset of nicotine exposure (Figure 2i-j). In the case of the FPE study, the body weight between nicotine-exposed and control mice became more similar after three weeks of exposure, and no clear changes were observed associated with nicotine cessation (Figure 2i). In contrast, the MPE study showed that the decrease in body weight between nicotine-exposed and control mice was maintained over time, albeit at varying degrees between exposure groups, and was reduced associated with nicotine cessation (Figure 2j).

Through the combined examination of water and food consumption and mouse body weight in FPE and MPE studies, we concluded that mice exposed to nicotine-laced water reduce water consumption, which may condition their feeding behavior and subsequently result in a decrease in body weight. Nicotine cessation also causes a rebound effect for water and food consumption, albeit to a lesser extent, and in body weight. Notably, female and male mice exhibit differences in their long-term acclimation to the consumption of nicotine-laced water, which may also influence food consumption.

### Effects of the parental exposure of female and male mice to nicotine and nicotine cessation

At the end of the exposure period in each study, we mated exposed mice with same-age mice of the opposite sex to engender their first descendant generation (F1). At 8-11 days of age, we counted and sexed F1 mice to determine litter size and sex bias. We compared litter size and the proportion of female offspring between each exposure group and the control group in each study using uMCW tests. Although a male sex bias was observed in litters for the three exposures when compared to controls in the FPE study, it was only statistically significant for LC (Figure 3a, b). Remarkably, despite the observation that tobacco smoking in humans reduced the likelihood of male births (43), a more comprehensive study of the influence of female smoking, considering parity, revealed that the offspring of primiparous smoking women tended to be male biased (44), as is the case in our studies.

**Figure 3.**
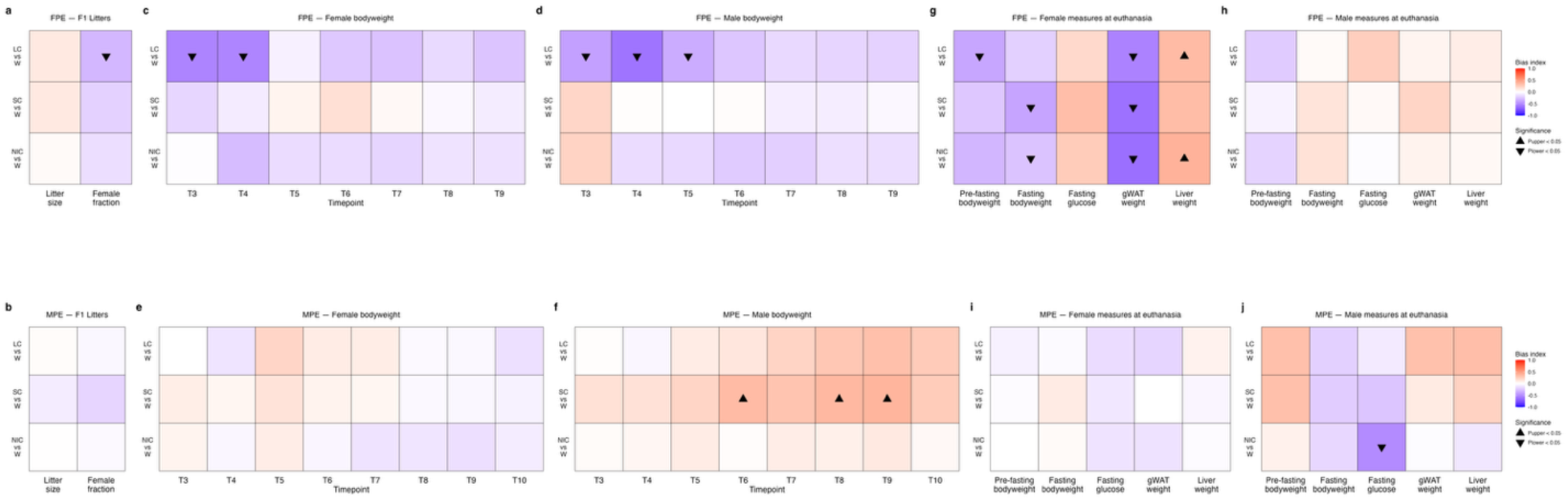
Effects of female and male preconception exposure of F0 female mice to nicotine and nicotine cessation on the metabolism of their F1 offspring. (**a, b**) Comparison of litter size (N) and sex ratio (female fraction) between exposure and control groups using uMCW tests for FPE (**a**) and MPE (**b**) studies. (**c-f**) Comparison of body weight normalized by age (g/days old) between exposure and control groups using uMCW tests for F1 females and males in FPE and MPE studies, separately. (**g, j**). Comparison of body weight normalized by age (g/days old), fasting body weight normalized by age (g/days old), fasting glucose normalized by body weight and age ((mg/dL)/g/days old), and tissue weight normalized by age (g/days old) between exposure and control groups for F1 females and males in FPE and MPE studies, separately. Sample size for each experimental group is >11 for litter size and female fraction analyses, and 10 for all other analyses. All comparisons were performed using uMCW tests, except for fasting body weight that were compared using mbMCW tests. FPE: female preconception exposure; MPE: male preconception exposure; mbMCW test: matched-bivariate measures Monte Carlo-Wilcoson test; uMCW test: unmatched measures Monte Carlo-Wilcoson test; BIs: bias index; W: water (negative control), LC: long cessation; SC: short cessation; NIC: nicotine.

To determine whether parental preconception exposure to nicotine and nicotine cessation resulted in metabolic perturbations in the F1 offspring of exposed mice, we selected one female-male sibling pair for ten litters per experimental group in each preconception study. We weighed the mice at weaning and weekly then after until they were at least nine weeks of age. Subsequently, we fasted mice for 12 hours, after which we determined fasting body weight and plasma glucose concentration and euthanized them to harvest cardiac blood samples, gonadal white adipose tissue (gWAT), and liver (Figure 1 and 3c-j). We used cardiac blood samples to quantify the plasma concentrations of twelve relevant metabolites (Figures 1 and 4).

**Figure 4.**
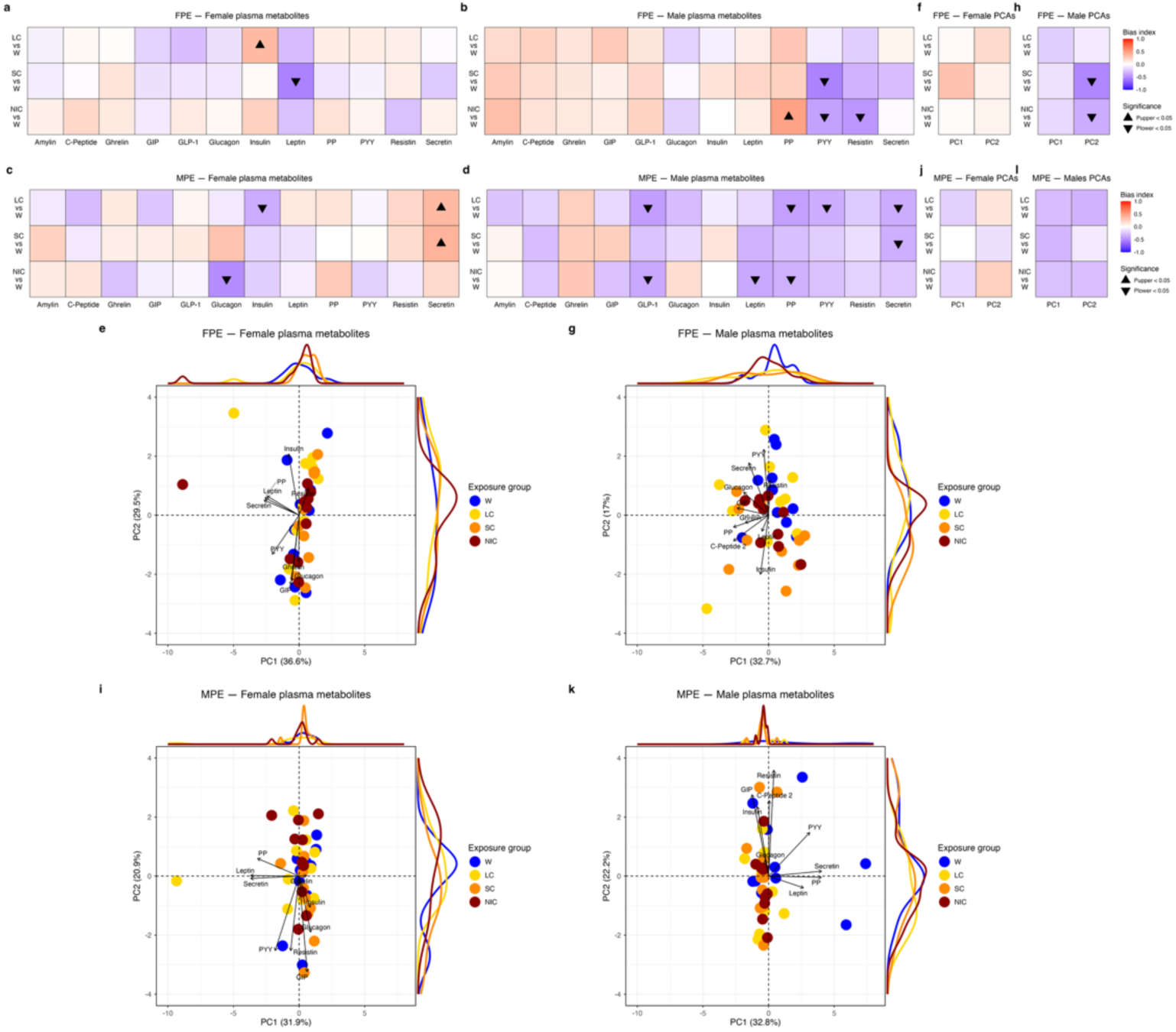
Effects of female and male preconception exposure of F0 female mice to nicotine and nicotine cessation on the plasma metabolites of their F1 offspring. (**a, d**) Comparison of plasma metabolite levels normalized by body weight and age ((pg/mL)/g/days old) between exposure and control groups using uMCW tests for F1 females and males in FPE and MPE studies, separately. (**e, g, i, k**) Principal component analyses (PCAs) of metabolite plasma levels normalized by body weight and age (pg/mL/g/days old) for all exposure and control groups for F1 females and males in FPE and MPE studies, separately. Principal component analyses (PCAs) of metabolite plasma levels normalized by body weight and age (pg/mL/g/days old) for all exposure and control groups of F1 females and males in FPE and MPE studies, separately. (**f, h, j, l**) Comparison of the distribution of mice along the two dimensions of PCA plots between exposure and control groups using uMCW tests for F1 females and males in FPE and MPE studies, separately. Sample size for each experimental group in all analyses is 10.

We assessed the significance of differences in body weight, fasting glucose, and tissue weight for each exposure *versus* control contrast using uMCW tests. We assessed the significance of differences in pre- and post-fasting body weights for each exposure versus control contrast using mbMCW tests.

Upon the joint analysis of weekly body weight, fasting body weight, fasting plasma glucose, and gWAT and liver weights, we observed sexual differences at two distinct levels (Figure 3c-j). Firstly, in both preconception studies, there were clear differences between F1 female and male offspring of mice exposed to nicotine and nicotine cessation. Secondly, the sexually dimorphic offspring effects of parental exposure to nicotine and nicotine cessation differed between FPE and MPE studies.

In the FPE study, F1 female mice exhibited a more pronounced effect caused by the maternal exposure to nicotine and nicotine cessation (Figure 3c-d, g-h). Although F1 weekly body weight tended to be lower in the LC and NIC groups compared to controls, and to a lesser extent in the SC group, the effects of FPE were more evident at euthanasia (Figure 3c, g). Body weight before and after a 12-hour fast, as well as gWAT weight, were lower in all three groups compared to controls (Figure 3g). Conversely, fasting glucose levels and liver weight were higher in the exposure groups compared to the control group (Figure 3g). While similar trends were observed for the three exposure *versus* control contrasts, their statistical significance varied (Figure 3c, g). Whether differences in tissue weights were significant in all three exposure *versus* control contrasts, body weight differences were only statistically significant in the LC *versus* W contrast, and fasting body weight differences were significant for the SC and NIC *versus* control contrasts (Figure 3c, g). In the case of F1 males, there was also some similarity in the trends for all three comparisons at euthanasia, but none of them were statistically significant and appeared considerably less biased than in females (Figure 3h). For F1 males, the only significant trends observed corresponded to lower body weights of LC mice compared to controls at weaning and the next two following weeks (Figure 3d).

In the MPE study, F1 male mice were the ones exhibiting a more pronounced effect caused by paternal nicotine exposure and cessation (Figure 3e-f, i-j). The body weight of LC, SC, and NIC mice tended to be higher than for control mice starting at 5 weeks of age (T5), being statistically significant only for LC and SC groups one or two weeks later, respectively (Figure 3f). At euthanasia, although there were some similarities in the trends for all exposure groups versus control contrasts for some traits such as body weight, fasting body weight, and fasting glucose, the only difference found significant was for fasting glucose in the NIC versus W contrast (Figure 3j).

We used plasma metabolite concentrations to conduct two distinct types of analyses. Initially, we compared data for each metabolite between each exposure *versus* control contrast for each F1 sex and preconception study using uMCW tests (Figure 4a-d). Subsequently, we assessed the similarities between all same-sex mice using principal component analysis (PCA) and data for all metabolites for each preconception study (Figure 4e-l).

In addition to the statistically significant differences observed for individual metabolite comparisons in nearly all but one exposure versus control contrasts, the uMCW test results reveal notable trends (Figure 4a-d). Although in F1 males for both studies, eight of the twelve metabolites demonstrated difference biases in the same direction for all contrasts, for the FPE study, most of the metabolites were elevated in the exposure groups when compared to controls, while for the MPE study, most of the metabolites were reduced (Figure 4b, d). The results of FPE and MPE F1 males also exhibited relevant trends in relation to the impact of nicotine cessation. In the FPE study, the results for PP, PYY, and resistin indicated that nicotine cessation mitigated the perturbations caused by female preconception exposure to nicotine, being the mitigation effect more pronounced in the longer cessation window (Figure 4b). Conversely, for the MPE study, the effect of nicotine cessation in mitigating the impact of nicotine exposure was not apparent (Figure 4d).

The PCA results for the FPE study were noteworthy. Visually, in FPE F1 males, W mice tend to preferentially occupy the upper right quadrant, while SC and NIC mice tend to be more distributed in the other three quadrants (Figure 4g). Remarkably, although LC mice are widespread, many of them appear to occupy a similar area of the graph as W mice (Figure 4g). The comparison of the distribution of data for the exposure and control groups in the two PCA dimensions separately using uMCW tests also supported the idea that the longer nicotine cessation window mitigated the effects of female preconception exposure to nicotine (Figure 4h). Upon integrating these findings with the individual analyses of the variations in plasma concentrations of individual metabolites (Figure 4b), it is plausible that the longest duration of nicotine cessation observed in this study may have a non-fully penetrant mitigating impact on the perturbations induced by the female preconception exposure to nicotine.

In the case of female mice in the FPE study, despite the distribution of LC, SC, and NIC mice appearing slightly different from that of W mice, particularly when considering PC1 (*x*-axis in Figure 4e), the separation of data for the exposure and control groups in the two PCA dimensions was not statistically significant when assessed using uMCW tests (Figure 4f). These results suggest that nicotine exposure may have a less significant impact on plasma metabolites of the F1 female offspring of females exposed to nicotine prior to conception, for whom the effect of nicotine cessation is not as evident. This conclusion is supported by the observed biases when assessing the differences between exposure and control groups for individual metabolites (Figure 4b).

The reason why the effects of nicotine and nicotine cessation exposures are more apparent in F1 females when analyzing physiological traits such as body weight, fasting glucose, or tissue weights, but more apparent in F1 males when analyzing the concentration of plasma metabolites is unclear.

In the context of MPE PCAs, a subtle distinction in the distribution of LC, SC, NIC, and W mice can only be discerned in the case of F1 males (Figure 4k-l). This observation aligns with the other findings of this study, which indicate comparable effects are observed across all exposure groups in F1 males, but not in F1 females (Figure 3e-f, i-j and 4c-d). Furthermore, these differences are more nuanced than those observed in the FPE study.

## Discussion

Numerous rodent studies have been conducted to elucidate the effects of nicotine exposure, encompassing the germ line of exposed mice, predominantly male, and their progeny (9,37,45–47). While a few studies have explored nicotine cessation in the germ line of exposed male mice (26,27,48), to the best of our knowledge, our research constitutes the first comparative analysis of the offspring effects of nicotine exposure and nicotine cessation in both sexes. Given that the primary objective of our research was to ascertain its validity, our studies are admittedly superficial. Despite the preliminary nature of our studies, some of our findings are noteworthy.

The patterns of water consumption we observed here demonstrated the anticipated nicotine aversion observed in mouse studies involving exposure to nicotine without odor or flavor masking (39,40,42). Notably, our studies revealed that male mice exhibited a gradual reduction in their aversion to nicotine-laced water over time, that was not detectable in female mice (Figure 2). While the aversion to consuming nicotine water and its subsequent decrease in prolonged exposures have been documented in male mice (42), similar studies have not been conducted in females possibly due to the primary focus on nicotine exposure in female mice during pregnancy and the utilization of additives that mask the odor or flavor of nicotine. Key areas for further investigation include the mechanisms underlying the sexual disparities observed in orally administered nicotine aversion, and whether these disparities correlate with human consumption patterns of unflavored and flavored nicotine products (49–51).

A highly relevant aspect of our research is that both female and male preconception exposures to nicotine and nicotine cessation elicited sexually dimorphic metabolic responses in their offspring (Figures 3 and 4). These findings are consistent with the possibility that offspring effects of preconception exposures are mediated by the alteration of the establishment of chromatin organization immediately after fertilization along the maternal-to-zygotic transition (MZT) (7).

First, it is well-established that diverse architectural elements of chromatin organization, such as the heterochromatin-euchromatin compartmentalization, are established during the initial two zygotic divisions (52–54). This critical moment in development is heavily reliant on the limited amount of material carried by gametes due to the transcriptional silencing of mature gametes and the early zygote (55). Any alteration of these germ line elements, for instance in response to environmental exposures, could subsequently lead to the establishment of perturbed chromatin organizations that could be propagated throughout development and across generations.

Second, during the final stage of spermatogenesis, the nuclear genome undergoes extensive compaction through the substitution of histones with protamines (56). Upon fertilization and prior to the first zygotic division, the protamines in the sperm chromosome are replaced by histones, a process that occurs entirely at the expense of the material deposited by the gametes (56). Given that the *Y* chromosome is predominantly heterochromatic in most species, it has been proposed that the *Y* chromosome may serve as a sink for heterochromatin-forming elements during periods of limited availability, such as the protamine-histone exchange of the sperm chromosomes immediately after fertilization (57,58). Consequently, it is anticipated that the establishment of chromatin organization on non-*Y* chromosomes will differ between females and males due to the presence of a *Y* or *X* chromosome among the sperm chromosomes. We found supporting evidence for this hypothesis by comparing gene expression stochastic variation between *Drosophila melanogaster XX* females and *XY* males, biological dispersal in species lacking *Y* chromosome homologs, and phenotypic heterogeneity in same-sex twin pairs (59,60).

The role of the mouse *Y* chromosome in the establishment of chromatin organization along the MZT remains uncertain. While the mouse *Y* chromosome presents a heterochromatic fraction smaller than in other species and several hundreds of highly expressed genes in the testis, most of these actively transcribed genes correspond to amplifications of three genes (61,62). Furthermore, it has been demonstrated that the heterochromatic fraction of the mouse *Y* chromosome can modulate epigenetic traits such as DNA methylation (63). Consequently, despite not fully conforming to the highly heterochromatic description of *Y* chromosomes in general, the mouse *Y* chromosome may still possess the potential to act as a modulator of chromatin organization during early embryogenesis.

Lastly, it is well-established that not only the maturation processes of oocytes and sperm are significantly distinct (14,15), but also that mature oocytes and sperm contribute differently to the material required for developmental processes occurring prior to the complete activation of the zygotic genome (64,65). Consequently, it is plausible that the same environmental exposure in females and males resulted in varying germ line perturbations, which in turn altered the establishment of chromatin organization in their offspring. This could potentially explain the diverse sexually dimorphic effects observed in the offspring of female and male mice exposed to nicotine-related preconception exposure in our studies (Figures 3 and 4).

In summary, the presence or absence of the *Y* chromosome in *XX* and *XY* zygotes might lead to a differential establishment of chromatin organization, which can subsequently be differentially impacted by the same perturbation of chromatin-forming elements carried by gametes. There may be other non-germline factors contributing to the offspring effects elicited by preconception exposures in our studies, such as perturbed physiologies during pregnancy of female mice previously exposed to nicotine or the possibility that male sperm can transport nicotine and cotinine that can directly affect the zygote (66,67). However, none of these factors would explain the sexual dimorphism we here detected in a straightforward manner. Further direct analyses are necessary to determine whether the environmental perturbation of germline elements, in conjunction with the effect of the presence or absence of the *Y* chromosome, is significantly associated with the offspring perturbation elicited by nicotine preconception exposures.

The results of the nicotine cessation windows further suggest specific research avenues to elucidate the potential germline effects of nicotine exposure. It is well-established that extended nicotine exposure to male mice disrupts their reproductive system and even the sperm methylome (26,48,68), which is minimized upon cessation windows encompassing entire cycles of sperm maturation (48). However, in our MPE study, we did not observe any effect consistent with an alleviating effect within our largest cessation window (Figures 3 and 4). It has been observed that nicotine exposure can alter the non-coding RNA load of sperm deposited while traversing the epididymis (45), and that RNAs play a crucial role in the establishment of the phase separation between heterochromatin and euchromatin (69–71). Therefore, although the mechanisms remain elusive, it is plausible that our findings indicate that nicotine might perturb the loading of non-coding RNAs in sperm which subsequently alter the establishment of chromatin organization in the offspring of exposure males, and that the cessation of nicotine exposure does not influence the nicotine-mediated perturbation of sperm non-coding RNAs.

In the FPE study, nicotine cessation appears to mitigate the effects of nicotine exposure on F1 male plasma metabolites, but not in F1 females (Figure 4). These findings suggest two potential avenues for further investigation. Firstly, the long cessation window, which spans one round of oocyte maturation, mitigates but does not fully reverse the effects of nicotine exposure. This implies that the germ line effect of nicotine may depend on the direct perturbation of nicotine or its metabolites during the oocyte maturation process (72), and that the cessation of nicotine exposure limits the effects. Secondly, the *Y* chromosome’s role as a sink for elements needed for the establishment of chromatin organization in the zygote becomes more apparent upon nicotine female preconception exposure. Nicotine exposure might effectively cause a reduction in these elements in oocytes, and nicotine cessation results in their restoration. In *XX* zygotes, the effects of nicotine are less evident due to the relatively minor reduction in these elements, making the effects of cessation less noticeable. In contrast, in *XY* zygotes, the *Y* chromosome captures a significant amount of these elements, resulting in even lower levels available for the establishment of chromatin organization in non-*Y* chromosomes than in *XX* zygotes. This hypothesis does not fully account for all of our findings. It remains unclear why F1 females, despite not showing clear effects in plasma metabolites, exhibit more pronounced effects on body and tissue weight at euthanasia not sensitive for the effect of cessation. Additionally, the viability of *XX* zygotes appears compromised, particularly in the long cessation window group (Figure 3a).

### Conclusion

Research conducted over the past several decades has demonstrated that the effects of environmental exposures are intricate and multifaceted. Consequently, multidisciplinary integrative frameworks are essential for a comprehensive understanding of these effects. For instance, the exposome concept has been defined as the integrated compilation of all physical, chemical, biological, and psychosocial factors, as well as their interactions, that influence biology and health, while exposomics is the transdisciplinary field dedicated to a thorough and discovery-driven comprehension of how the exposome impacts biology and health (73–75). Furthermore, the complexity of environmental exposures and their effects is manifested in the need of up to twelve key characteristics to effectively define metabolism-disrupting agents (11).

Our studies, albeit intentionally simple, were designed to leverage biological complexity—represented here by the different gamete dynamics in females and males—to assess whether complex exposures that include cessation prior to conception lead to offspring effects that can be mitigated by that cessation. Despite the interpretive challenges in explaining all of our results, nicotine and nicotine cessation, examined through the lens of gamete development, offer a useful approach for understanding the mechanisms underlying nicotine’s multigenerational effects.

We believe this research carries both fundamental and translational significance. It will help elucidate the mechanisms underlying the multigenerational environmental determination of disease predisposition, and, by focusing on preconception exposures, it can guide the implementation of cessation strategies that mitigate or prevent such adverse effects. More comprehensive characterization of both direct and offspring effects remains necessary to fully assess the multigenerational impact of nicotine consumption. Establishing a causal link between the two is crucial for identifying individuals at heightened risk of transmitting these effects to their offspring, as is the development of more precise prevention strategies.

## Data and code availability

The data and code necessary to replicate the analyses presented in this study will be publicly distributed upon peer-reviewed publication.

## Ethics

These procedures were reviewed and approved by the UCSC Institutional Animal Care and Use Committee (UCSC IACUC) as part of the animal protocol Diazc2204.

## Author contributions

AP: Investigation, Writing — review & editing. RC-G: Conceptualization, Funding acquisition, Investigation, Resources, Supervision, Writing — review & editing. CD-C: Conceptualization, Data curation, Formal analysis, Funding acquisition, Investigation, Resources, Software, Supervision, Validation, Visualization, Writing — original draft, Writing — review & editing

## Acknowledgments

We express our profound appreciation to Stephanie R. Aguiar, Daniel Davis, Truman Natividad, Maya Skinner, Bailey Thompson, and Ewan Whittaker Walker for their invaluable contributions to the mouse dissections conducted for this research.

## Generative AI statement

The author(s) declared that generative AI was not used in the creation of this manuscript.

